# Cell-free transcriptional regulation via nucleic-acid-based transcription factors

**DOI:** 10.1101/644021

**Authors:** Leo Y.T. Chou, William M. Shih

## Abstract

Cells execute complex transcriptional programs by deploying distinct protein regulatory assemblies that interact with cis-regulatory elements throughout the genome. Using concepts from DNA nanotechnology, we synthetically recapitulated this feature in cell-free gene networks actuated by T7 RNA polymerase (RNAP). Our approach involves engineering nucleic-acid hybridization interactions between a T7 RNAP site-specifically functionalized with single-stranded DNA (ssDNA), templates displaying cis-regulatory ssDNA domains, and auxiliary nucleic-acid assemblies acting as artificial transcription factors (TFs). By relying on nucleic-acid hybridization, de novo regulatory assemblies can be computationally designed to emulate features of protein-based TFs, such as cooperativity and combinatorial binding, while offering unique advantages such as programmability, chemical stability, and scalability. We illustrate the use of nucleic-acid TFs to implement transcriptional logic, cascading, feedback, and multiplexing. This framework will enable rapid prototyping of increasingly complex in vitro genetic devices for applications such as portable diagnostics, bio-analysis, and the design of adaptive materials.

## Introduction

Living cells use information encoded in biochemical circuits to make complex decisions and perform sophisticated tasks. Inspired by their rich functionality, synthetic gene circuits are currently being developed to model biology and engineer organisms for various applications.^1,2^ Recently, there has also been increasing interest to create gene circuits that operate in vitro using reconstituted molecular components. Compared to cellular devices, these cell-free ones are more portable, accessible, and robust. These advantages are now being explored for applications such as point-of-care diagnostics^3^, artificial cells^4–6^, expression of toxic products^7^, screening^8^, and even for educational purposes^9^. Yet as with cellular devices, scaling up the complexity of synthetic gene circuits requires a large toolbox of regulatory elements that can wire up genetic elements without introducing cross-talk. In living cells, this circuit wiring is achieved via interactions between TFs with cis-regulatory elements distributed throughout the genome. The molecular properties of TFs enable sophisticated self-assembly-mediated regulatory behaviors, including recognition of specific promoters^10^, recruitment of co-regulatory units, signal integration via multi-component assembly^11^, and even physical alteration of genome structure^12^. Engineering these regulatory behaviors has been a rate-limiting step in the design of synthetic gene circuits.

In contrast to proteins, nucleic-acid-based regulatory elements offer a solution for programmable gene regulation by relying on Watson-Crick hybridization for predictable self-assembly, and by taking advantage of the sophisticated software tools that are available to predict nucleic-acid interactions. Recently, several synthetic, programmable RNA-based regulatory devices have been developed, such as toehold switches^13^ and small transcriptional activating RNAs (STARs)^14^, which either regulate transcription elongation or the translation of mRNA into proteins. In contrast, fewer programmable mechanisms exist for regulation at the level of transcription initiation. Pioneering efforts to engineer in vitro transcriptional networks using nucleic acids have largely focused on manipulating the interactions of T7 RNAP with its promoter by either making the promoter single-stranded (i.e. inactive) or double-stranded (i.e. active).^15^ While this strategy has enabled the construction of in vitro circuits with interesting dynamics^16,17^, its scalability is restricted by the T7 promoter sequence. Here we describe a new regulatory architecture for in vitro transcriptional regulation that alleviates this constraint. This architecture supports the use of arbitrary sequences of DNA or RNA as inputs to produce arbitrary RNA-based outputs, making the transcriptional network modular and composable (Fig. 1), which lends itself to standardization, abstraction, and scaling.

**Figure 1.**
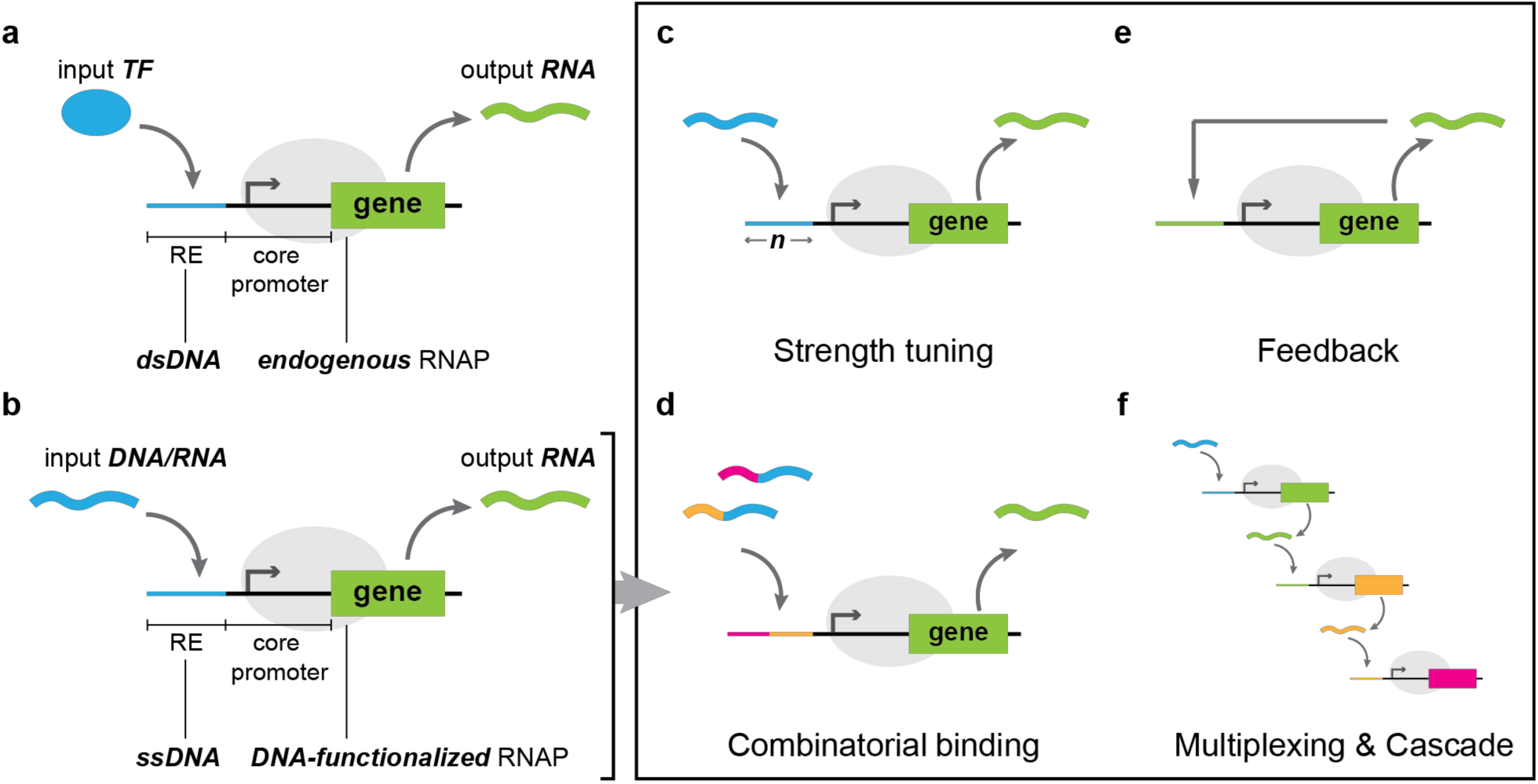
Overview of nucleic-acid-regulated transcription. **a**, General schematic of endogenous transcriptional regulation. Protein-based transcription factors (TFs) bind to regulatory element (RE) upstream of the core promoter region to either enhance or suppress gene expression. **b**, Nucleic-acid-based regulatory architecture developed in this study. Instead of using protein-based transcription factors, DNA/RNA regulatory assemblies are engineered to interact with ssDNA *cis-*regulatory elements via sequence-programmable hybridization for local enhancement or suppression of the activity of a DNA-functionalized T7 RNAP. Since both input and output of the gene are in the form of nucleic acids, and there are no sequence constraints, this mechanism of gene regulation is modular and composable, allowing for the rational design of a number of behaviors including the following: **c**, programmable transcriptional activation with tunable strengths via the length of the RE domain, *n*, **d**, combinatorial and cooperative activation, **e**, feedback, and **f**, multiplexing and cascading.

## Results

### Programmable transcription using a “caged” DNA-functionalized T7 RNAP

The wild-type T7 RNAP lacks regulatory mechanisms beyond its ability to recognize and bind to its 17-bp promoter. To expand on its regulatory capacity, we created a ssDNA-functionalized T7 RNAP by covalently coupling a 21-nt single-stranded DNA to a recombinant T7 RNAP bearing an N-terminal SNAP-tag (Fig. S1) Hybridization of a synthetic duplex to this ssDNA-tag yielded a “caged” T7 RNAP whose activity could be controlled in programmable fashion via DNA strand displacement (Fig. 2a).^18^ The cage duplex encodes a truncated T7 promoter (P_T7, ΔGGG_) that retains affinity for the RNAP active site but lacks the initiation sequence (e.g. “GGG”) necessary for RNAP to undergo conformational switching into its transcriptionally competent, elongation state (Fig. 2b).^19–21^ Mechanistically, the cage works by trapping the RNAP in its initiation conformation and inhibiting its ability to associate with other substrates in solution (Fig. S2). In addition, the cage is highly efficient by virtue of being in close proximity to the RNAP. The activity of the RNAP is fully recovered upon a simple strand-displacement operation. For example, a template can be programed to display a complementary ssDNA “operator” domain that invades and displaces the cage duplex from the RNAP-cage complex (Fig. 2c). By placing the operator upstream of the promoter of the template, the uncaged RNAP is primed to initiate transcription of the downstream gene.

**Figure 2.**
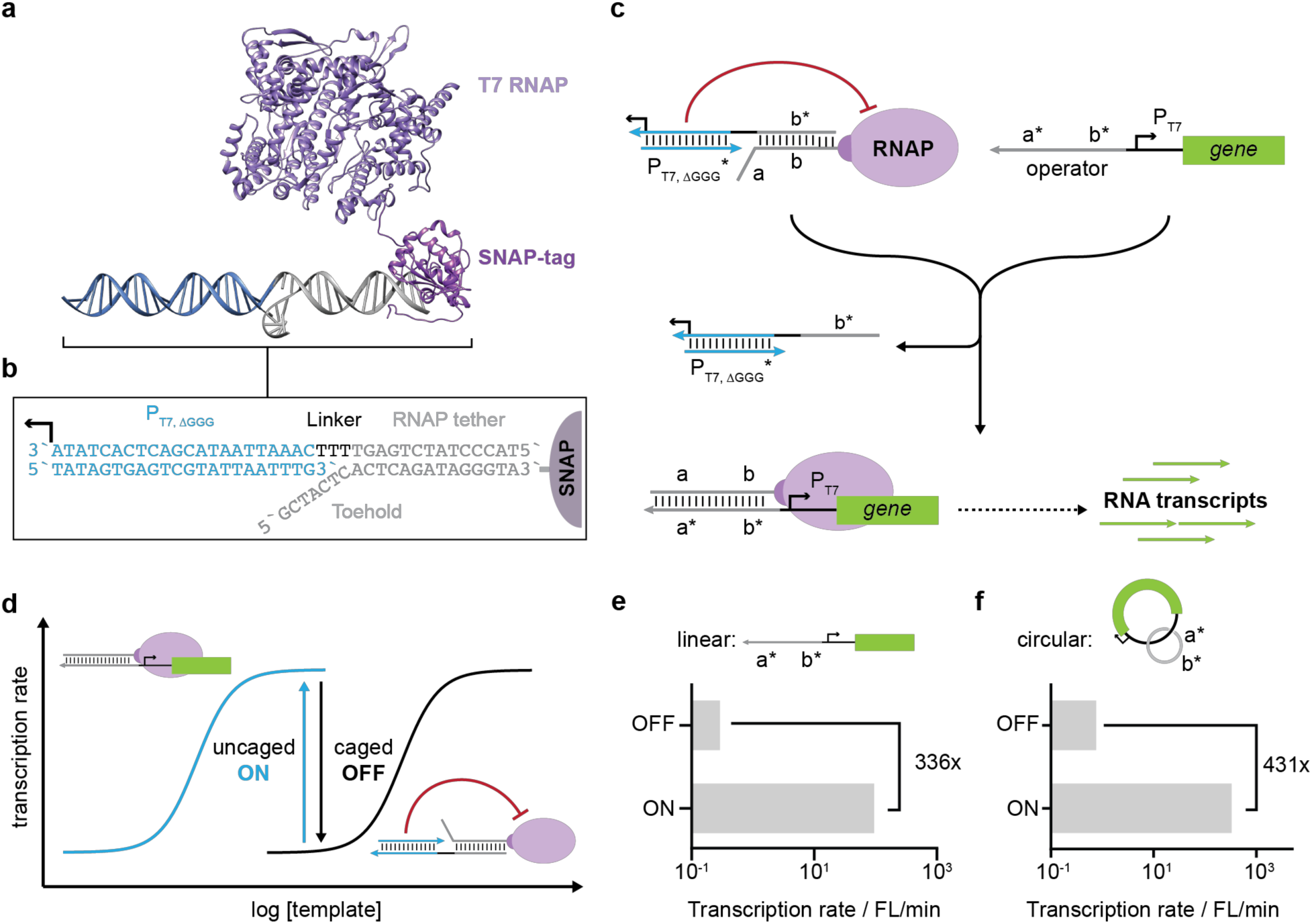
Transcriptional activation for DNA-functionalized T7 RNAP system. **a**, Schematic of the “caged” T7 RNAP used in this study. Recombinantly expressed T7 RNAP bearing N-terminal SNAP-tag is covalently conjugated to a 21-nt ssDNA (grey strand). Hybridization of a “cage” duplex to this ssDNA-tag yields an RNAP whose activity is activatable via programmed DNA strand-displacement. **b**, The cage duplex encodes a truncated T7 promoter (blue domain), which fails to induce transcription but nevertheless occupies the active site of the RNAP and prevents its association with other DNA templates. The duplex can be removed via strand displacement mediated by a 7-nt toehold positioned at the 5’ end of the ssDNA-tag. **c**, Example schematic of a strand-displacement reaction for RNAP uncaging and loading onto a gene-of-interest. **d**, Relationship between enzyme kinetics and cage state. Hybridization of cage duplex to RNAP introduces a locally bound competitor for template binding, resulting in a large shift in RNAP activity between the caged vs. uncaged states at most concentrations of template. **e**,**f**, Transcription velocity of the RNAP, here monitored as the rate of production of a fluorescent RNA aptamer per unit time, in the caged (i.e. OFF) vs. uncaged (i.e. ON) states, measured on either linear (e) or circular templates (f).

From an enzyme-kinetics perspective, the cage acts as a highly efficient, intramolecular competitor when bound to the RNAP via its ssDNA-tag. The transition from RNAP-cage complex to RNAP-template complex reverses this situation, resulting in a large change in transcriptional velocity between the caged (i.e. OFF) and the uncaged (i.e. ON) states (Fig. 2d). We identified parameters that affect the dynamic range between these two states by monitoring the production of a fluorescent RNA aptamer (e.g. Broccoli) under both conditions. We found that Broccoli expression in the ON state scaled linearly with the length of the ssDNA domain on the template by stabilizing RNAP binding (Fig. S3). On the other hand, Broccoli expression in the OFF state was determined by caging efficiency, which varied as a function of cage sequence, concentration, tether stability, and buffer ionic strength (Fig. S4). Optimizing these conditions reduced OFF-state expression to near background levels, resulting in 336-fold gene activation (Fig. 2e, see also Fig. S5). As a more stringent test, we repeated this experiment using a circular DNA template catenated with another circular ssDNA to which RNAP can localize and produce RNA continuously via rolling-circle transcription (Fig. S5). As with the linear template, we achieved >400-fold gene activation by inhibiting OFF-state transcription (Fig. 2f, see also Fig. S6). Together, these results illustrate programmable activation of our ssDNA-functionalized T7 RNAP using nucleic-acid hybridization.

### Nucleic-acid structures as synthetic transcription factors

To introduce additional regulatory mechanisms, we programmed the RNAP to co-localize with templates via auxiliary nucleic-acid structures serving as artificial TFs. As two examples, we designed nucleic-acid repressors that respectively emulate the inducible (e.g. *lac*) and repressible (e.g. *trp*) gene systems in *E. coli* (Fig. 3a,d). These systems were picked because they have been the workhorses of synthetic gene networks, and therefore, the construction of functional nucleic-acid analogs may similarly provide the basis for building more complex cell-free circuits. In the *lac* system, the repressor protein binds to the operator domain of the gene to block the RNAP from engaging with the promoter. Repression is allosterically alleviated by effector molecules (e.g., allolactose) binding to the repressor (Fig. 3a). Our mimic of the *lac* repressor consists of a linear strand that blocks the DNA-functionalized T7 RNAP from binding to the ssDNA operator on the template. De-repression occurs when the effector strand removes the repressor strand from the template via toehold-mediated strand displacement, allowing template-mediated uncaging of RNAP (Fig. 3b). Figure 3c shows the dose-response of this scheme as a function of effector-strand concentration. The response is much sharper compared to typical *lac-*allolactose systems, reflective of the stronger binding energetics between nucleic-acid strands. Notably, the dynamic range of the dose-response is systematically tunable via sequence design. For example, the inset in Figure 3c shows how the response for a given effector concentration decreases as the length of its hybridizing domain decreases (see also Fig. S7). In contrast to the lac system, the *trp* repressor consists of effector and repressor molecules associating cooperatively to suppress gene expression, which is the logical equivalent of a digital NAND gate (Fig. 3d). We recapitulated this logic by designing two nucleic-acid strands that associate to form a four-way junction (4WJ) with the regulatory operator domain on the template (Fig. 3e&f). Inset in Figure 3f shows how the repression changes as a function of deletions in the hybridization domain between the effector and repressor strands. In addition, a more graded dose-response can be achieved by replacing the 4WJ motif in the *trp* mimic with a three-way junction (3WJ, Fig. S8), reflecting the known weaker binding energetics of 3WJ compared to 4WJs.^22^ These examples highlight how DNA nanotechnology design principles can be used for precise engineering of gene expression profiles.

**Figure 3.**
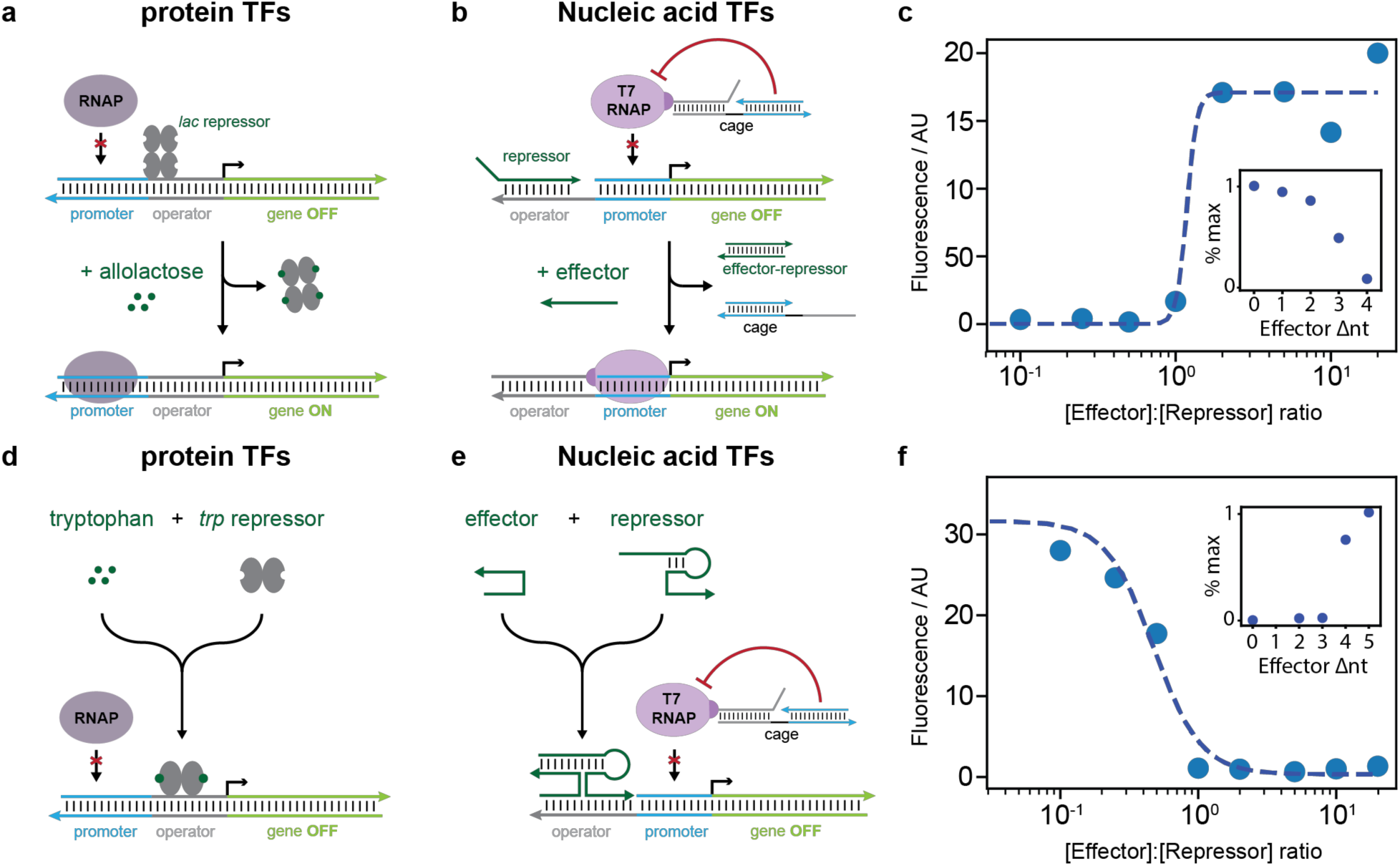
Synthetic recapitulation of endogenous gene-regulatory architectures. **a**, Schematic of a *lac-*inducible gene in *E. coli*. Binding of the *lac-*repressor protein to the operator on the DNA template prevents RNAP binding to the promoter, resulting in gene repression. Repression is alleviated upon allosteric binding of allolactose to the *lac-*repressor protein, which turns on gene expression. **b**, Nucleic-acid mimic of the *lac* system. A linear ssDNA acts as the repressor by binding to the operator region of the DNA template, masking the toehold required for template-mediated RNAP uncaging. An effector strand displaces the repressor from the template to trigger gene expression. **c**, Dose-response of the nucleic-acid *lac* mimic as a function of the effector-to-repressor ratio. Inset shows how the maximum response varies as a function of decreasing hybridization length between the effector and repressor from 0 to 4 nt. **d**, The *trp* repressible gene. Binding of tryptophan to the *trp* repressor allosterically strengthens its affinity for the operator, resulting in gene suppression. In the absence of the tryptophan, the *trp* repressor is unable to suppress gene expression. **e**, Nucleic-acid mimic of the *trp* system. The effector and repressor strands assemble to form a four-way junction (4WJ) on the template, thereby preventing template-mediated RNAP uncaging. **f**, Dose-response of the nucleic-acid *trp* mimic as a function of the effector-to-repressor ratio. Inset shows how the fluorescence signal varies as a function of the length of the hybridizing region between effector and repressor for a given effector concentration.

### Feedback, cascading, and transcriptional multiplexing

In addition to using DNA as transcriptional regulators, our system can also be regulated using RNA, such as using nascent transcripts to execute feedback and/or cascading. As an example, we created an auto-inhibitory circuit by constructing a gene that encodes its own repressible effector molecule (Fig. 4a). Expression of this gene produced RNA molecules that combine with free-floating DNA repressors that cooperatively inhibit its own transcription (Fig. 4b). As another example, we constructed a two-step cascade with auto-catalytic feedback (Fig. 4c). The first step in the cascade is a constitutively active template that produces effectors to activate the second template. The second template is auto-catalytic because it also produces its own effectors, but it is initially inhibited by excess DNA repressors. The response of the system is an exponential increase in gene expression, in this case of a fluorescent RNA aptamer, at different time points that is determined by the initial concentration of the DNA repressor (Fig. 4d & S9). Together, these results demonstrate how both DNA and RNA can be used as TFs to execute transcriptional logic and feedback. In addition, this RNA-in, RNA-out regulatory mechanism makes it relatively straightforward to design genetic elements and assemble them into circuits (e.g. composable).

**Figure 4.**
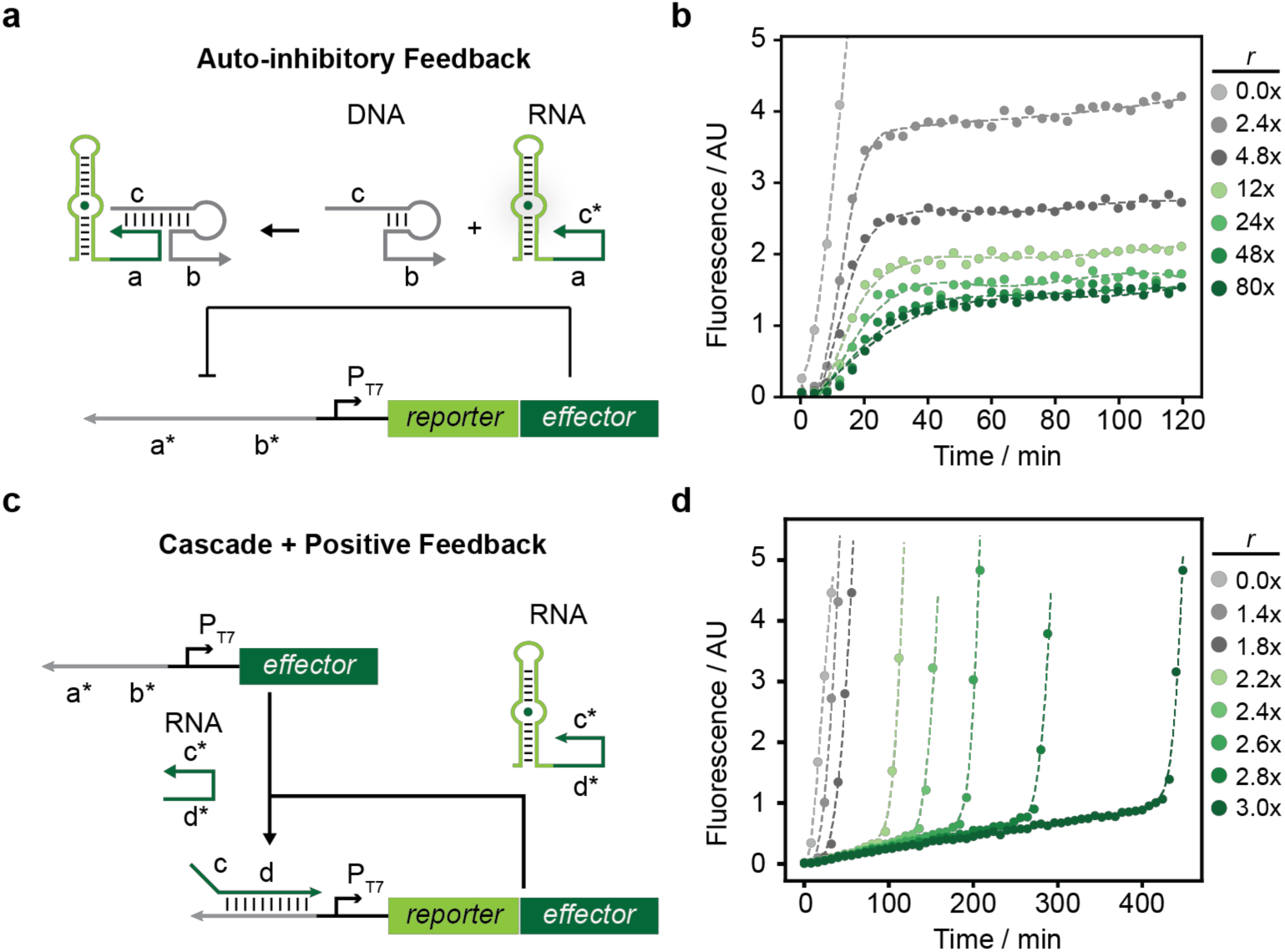
Implementation of negative and positive feedback. **a**, Schematic of an auto-inhibitory feedback loop. The DNA template encodes a fluorescent RNA aptamer (reporter) attached at its 3’-end to an effector sequence. Upon transcription, this RNA molecule assembles with free DNA repressors in solution for cooperative suppression of its own production. **b**, Kinetics of the auto-inhibitory response as a function of the initial repressor-to-template ratio, *r*. **c**, Schematic of a two-step cascade driving an auto-catalytic feedback loop. The first template in the cascade is constitutively active and produces an effector sequence that removes the DNA repressor initially bound to the second template in the cascade. The second template encodes an auto-catalytic RNA molecule consisting of a fluorescent RNA aptamer (reporter) attached at its 3’-end to the same effector which, when transcribed, alleviates its own inhibition by the repressors. **d**, Kinetics of the auto-catalytic response as a function of the initial repressor-to-template ratio, *r*.

Another unique advantage of nucleic-acid-based TFs is their scalability. Because DNA hybridization relies on Watson-Crick base pairing, many instances of the same molecular motif can be created by assigning unique sequence choices for each logical domain. We demonstrate this by operating twelve instances of a regulatory motif in a multiplexed format. The regulatory motif we are using is shown in Figure 5a, where a gene is activated upon docking of a pair of nucleic acid TFs, denoted TF_A_ and TF_B_, equivalent to a digital AND gate. We designed a set of twelve orthogonal templates based on this architecture, each encoding for a different RNA transcript “barcode”, and each regulated by an orthogonal pair of TFs (Table S9). To test multiplexed gene activation, we combined the templates into a pool and performed twelve independent *in vitro* transcription reactions using this template pool, each activated using one pair of TFs. We then assayed for the identity of the RNA barcode transcribed using a set of molecular beacons each specific for one RNA barcode (Fig. 5b). As a first-stage verification, we visualized the transcription reactions via denaturing PAGE (Fig. 5c). Here we observed RNA production for all twelve TF pairs added to the template pool (“ALL”, Fig. 5b). With the exception of TF pairs 4 and 11, the expression levels varied by less than 2-fold across all designs (Fig. 5c, bottom graph). More importantly, repeating this experiment with the target template removed from the pool resulted in no detectable gene expression, suggesting that the RNA production we trigger using the TF pair is specific to its cognate template (“LOO”, Fig. 5b). We further validated this design by using the RNA transcripts to activate molecular beacons (Fig. 5d). The results demonstrate highly specific molecular-beacon activation with minimal cross-talk, consistent with the notion of orthogonal transcriptional activation in a multiplexed format. These results demonstrate the possibility of rapidly prototyping de novo regulatory elements for multiplexed operation.

**Figure 5.**
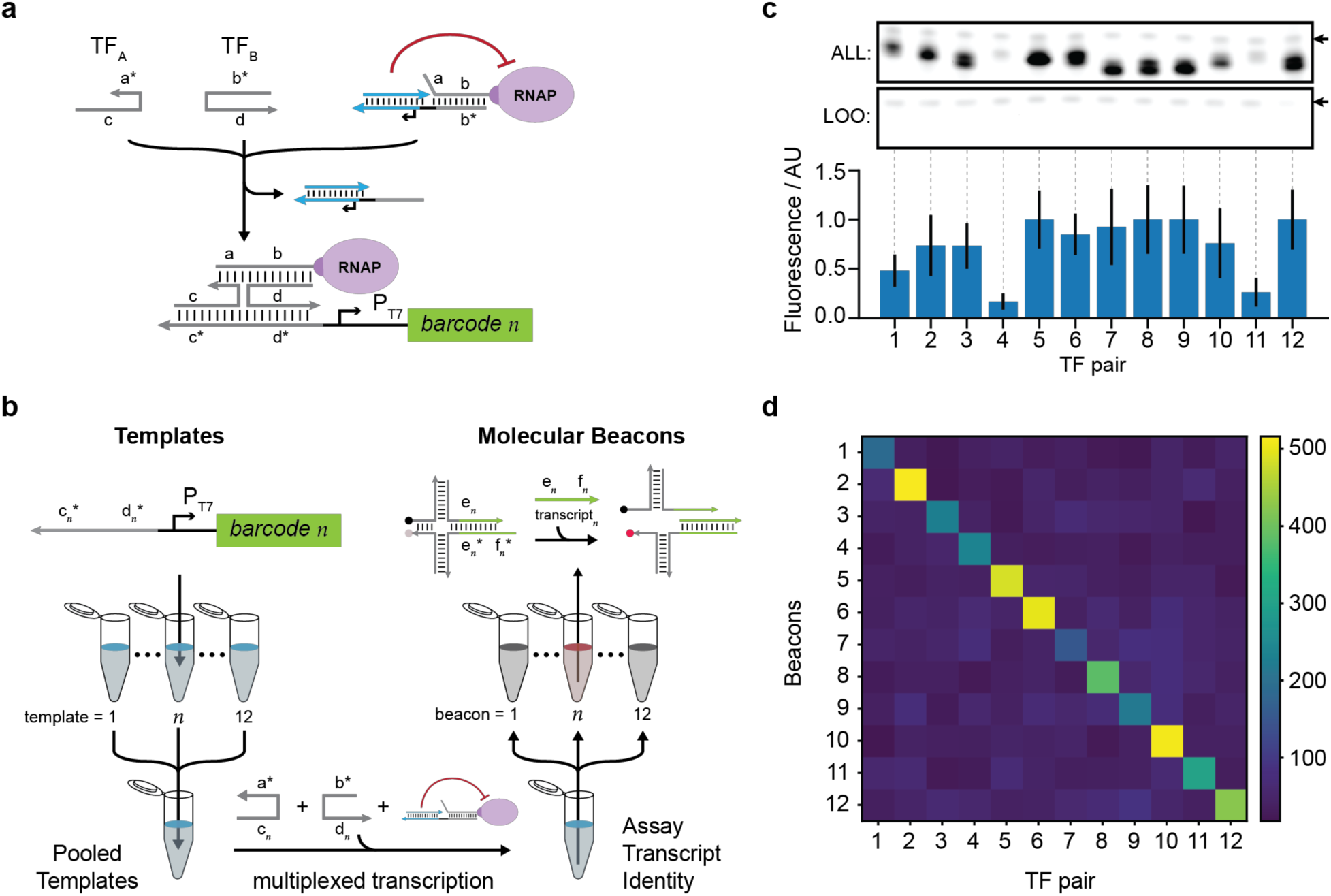
Implementation of transcriptional multiplexing. **a**, Schematic of the regulatory architecture of the genetic templates used for testing multiplexed transcription. Each template *n* encodes a unique RNA barcode *n* whose transcription is activated upon the binding of a pair of nucleic acids, denoted TF_A_ and TF_B_, to the template operator. **b**, Schematic of the experiment used for multiplexed transcription and RNA identification. Twelve templates each encoding a different RNA barcode and under the regulation by the architecture shown in **a** are combined into a pool. This pool is used to set up twelve independent in vitro transcription reactions each performed using one TF pair. The identity of the RNA transcribed from each reaction is verified using a set of twelve molecular beacons specific for each RNA barcode. **c**, TBE-Urea PAGE showing RNA produced from the twelve independent multiplexed in vitro transcription reactions. “ALL”: in vitro transcriptions using a template pool containing all 12 templates. “LOO”: in vitro transcriptions using a “leave-one-out” pool, in which the cognate template corresponding to the TF pair added was removed from the pool. Arrow in the gels points to the cage duplex, here used as a loading control to normalize signals across samples. Bottom: Quantification of RNA produced from the transcription reactions from the “ALL” template pool normalized to the highest level. Error bars denote standard deviation from three independent experiments. **d**, Activation of molecular beacons by RNA produced from each multiplexed transcription reaction displayed as an activation matrix. Signals across matrix diagonal represent specific activation while off-diagonals indicate nonspecific activation.

## Discussion

Gene-regulatory networks play an integral role in both native and synthetic gene circuits. By defining the logical connectivity between genes, they enable gene circuits to process and respond to complex environmental signals. The function of gene-regulatory networks derives from faithful self-assembly-based molecular recognition between regulatory components. A central effort in the field of synthetic biology has been to engineer these components in order to construct circuits with increasing computational power. Developing de novo protein-based TFs for transcriptional regulation has proven especially challenging due to the complexity of designing TF-TF and TF-promoter interfaces. In this study, we circumvented this complexity by developing nucleic-acid-based TFs for transcriptional regulation of in vitro synthetic gene circuits. This framework uses synthetic nucleic-acid assemblies to program the association and subsequent activity of an ssDNA-functionalized T7 RNAP with its DNA templates, analogous to how endogenous TFs interact with cis-regulatory elements to recruit RNAP to genomic start sites. By replacing protein-DNA interactions with nucleic-acid hybridization, we were able to rapidly prototype gene-regulatory behaviors and create large numbers of orthogonal regulatory elements. While nucleic acids have been previously used to control T7 RNAP activity^15,23^, these studies have largely focused on switching the state of the promoter between ssDNA versus dsDNA, which poses inherent sequence and structural constraints. By alleviating these constraints, our framework renders the design of such nucleic-acid regulatory assemblies modular and composable, which lends itself to abstraction, standardization, and scaling. These advantages will make it easier to construct increasingly sophisticated cell-free genetic circuits for various applications.

We propose a number of future directions for further advancing this technology. The first is to realize large libraries of standardized regulatory components. In contrast to proteins, such DNA-based regulatory components should be easier to design, tune, and characterize. As proof-of-concept in this study, we developed a panel of twelve orthogonal nucleic-acid transcription factors (e.g., Fig. 5). Future studies could expand on the size of this panel, as well as panels of other regulatory motifs, to hundreds of standardized parts, which will broadly support the design of more complex circuits. Second, we envision future iterations of this technology to interface even more closely with DNA-based computing technologies. Systems of synthetic oligonucleotides have been successfully designed as switches, amplifiers, logic gates, and oscillators.^24–27^ By programming these circuits to produce specific TF sequences as outputs, they can function as embedded controllers for programming gene-expression dynamics under our framework. This use of nucleic-acid computing for the active, on-demand synthesis of functional RNAs could find applications in biological analysis, directed evolution, and molecular information processing. Third, we foresee ample opportunities for synthetic recapitulation of native gene-regulatory mechanisms using DNA nanotechnology. In this study, we created nucleic-acid inducible and repressible genes by mimicking the structure of a prokaryotic operon (e.g., Fig. 3). In the future, co-regulated gene clusters can be envisioned by designing nucleic-acid scaffolds that mediate higher-order organization of genes and TFs, analogous to the actions of endogenous long-noncoding RNAs^28^, or by organizing genes into artificial DNA nanostructures that can reconfigure in analogous fashion to chromatin reorganization^29^. These efforts will enable more sophisticated levels of synthetic gene regulation. Finally, yet still more refined gene regulation can be explored by merging our work with those operating at the post-transcriptional levels^13,14^ and with methods based on spatial patterning and compartmentalization^6^.

Applications of cell-free synthetic gene circuits are now beginning to emerge, such as portable diagnostics, distributed bio-manufacturing, and therapeutic artificial-cells.^3,30,31^ Regulating gene-expression dynamics is desirable in these applications in order to focus a finite amount of energy and resources towards manufacturing the right product at the right time. Compared to protein-based gene-regulatory frameworks, the nucleic-acid-based regulatory framework presented in this study offers a number of functional advantages for synthetic gene-regulation. First, nucleic-acid regulatory elements consume less resources and can potentially accelerate circuit-response time because their production does not involve the translation machinery. Second, for devices designed for portability, nucleic acids have favorable storage and distribution characteristics compared to proteins. Finally, for circuits being developed for point-of-care diagnostics, nucleic-acid regulated circuits can directly interface with DNA and RNA molecules extracted from physiological fluids, or else with small molecules and proteins via the use of aptamers or DNA-encoded affinity agents. An intrinsic constraint of our approach is the need to synthesize polymerase-DNA conjugates and genes containing ssDNA domains. Nonetheless, we believe these efforts will be scalable^32^ and offset by the programmability and gains in performance offered by nucleic-acid-based gene regulation.

## Acknowledgements

L.Y.T.C. acknowledges the Banting Postdoctoral Fellowship for generous support. This work was funded by support from NSF Expeditions CCF-1317291 and the Wyss Institute at Harvard Core Faculty Award to W.M.S. The authors would like to thank Dr. Rasmus Sørensen and Dr. Jaeseung Hahn for helpful discussions.

## Author Contributions

L.Y.T.C. conceived the project, planned and executed the experiments, and co-wrote the manuscript. W.M.S planned experiments, co-wrote the manuscript, and supervised the project.

